# IDENTIFICATION OF KEY GENES ASSOCIATED WITH PCOS (POLY CYSTIC OVARIAN SYNDROME) AND ENDOMETRIAL/OVARIAN CANCER THROUGH BIOINFORMATICS

**DOI:** 10.1101/2024.01.23.576969

**Authors:** Karishma Raulo, Sahar Qazi

**Affiliations:** Quick IsCool, Aitele Research LLP, Bihar, India

**Keywords:** PCOS, ENDOMETRIAL CANCER, OVARIAN CANCER, BIOINFORMATICS

## Abstract

**Background:** PCOS, a common endocrine disorder, is linked to increased risks of endometrial cancer (EC) and ovarian cancer (OC). Our study utilizes bioinformatics analysis to identify shared gene signatures and elucidate biological processes between EC/OC and PCOS.

**Materials and Methods:** Gene expression profiles for PCOS (GSE199225), endometrial cancer (GSE215413), and ovarian cancer (GSE174670) were obtained from the GEO database. Hub genes were identified through functional enrichment analysis and protein-protein interaction. Drug identification analyses were employed to find drugs targeting the hub genes.

**Result:** Key hub genes linking PCOS and EC includes RECQL4, RAD54L, ATR, CHTF18, WDHD1, CDT1, PLK1, PKMYT1, RAD18, and RPL3; for PCOS and OC, they include HMOX1, TXNRD1, NQO1, GCLC, GSTP1, PRDX1, SOD1, GPX3, BOP1, and BYSL.

Gene Ontology analysis revealed DNA Metabolic Processes in PCOS and EC, while in PCOS and OC, it identified the Removal of Superoxide Radicals. KEGG Pathway analysis highlighted Cell cycle in PCOS and EC and Hepatocellular carcinoma in PCOS and OC. Potential drugs for PCOS and EC include quercetin, calcitriol, and testosterone; for PCOS and OC, eugenol and 1-chloro-2,4-dinitrobenzene are identified.

**Conclusion:** These findings offer insights into potential therapeutic targets and pathways linking PCOS with EC and OC, enhancing our understanding of the molecular mechanisms involved in these associations.

## INTRODUCTION

Polycystic Ovary Syndrome (PCOS) impacts around 5 to 10% of women in their reproductive years, emerging as the leading cause of anovulation among those facing infertility. Its onset is gradual, presenting a clinical spectrum that encompasses the classic triad of PCOS symptoms: hyperandrogenism, menstrual irregularities, and the presence of polycystic ovaries. [1]

A polycystic ovary detected by ultrasound, infertility, acne, amenorrhea or oligomenorrhea, hirsutism, insulin resistance, obesity, and hyperandrogenism can all be indicators of PCOS.[2] In addition to reproductive irregularities, a variety of metabolic illnesses, including hypertension, hepatic steatosis, glucose intolerance, dyslipidemia, and type II diabetes, are significantly linked to PCOS. In the development of PCOS, hyaluronic acid (HA) plays a crucial role in fostering ovulatory dysfunction. It induces abnormalities in lipid metabolism, encourages hyperinsulinemia and insulin resistance, disrupts the LH-to-FSH ratio, and raises the frequency and intensity of LH and GnRH pulse secretion. Studies have suggested that these factors possess the capacity to directly stimulate the proliferation of cancer cells.[3]

Studies have established a connection between PCOS and cancers affecting the endometrium, ovaries, kidneys, hematological system, and pancreas. Progress in comprehending the molecular processes involved in PCOS has facilitated these discoveries.[4]

Endometrial cancer (EC) is a major gynaecological concern, prevalent in the Western world, and a significant cause of mortality. It affects the inner lining of the uterus, with a globally increasing incidence. Meta-analyses highlight polycystic ovary syndrome (PCOS) as a notable risk factor, with women having PCOS being three times more likely to develop EC. The risk is significantly higher for women under 54 compared to older individuals [5]. The primary hypotheses seeking to elucidate the connection between PCOS and EC involve elevated estrogen levels, hyperinsulinemia, progesterone resistance, and decreased apoptosis in women. These factors contribute to the development of endometrial hyperplasia, ultimately leading to the onset of EC. [6]

Ovarian cancer currently ranks as the fifth leading cause of cancer-related deaths among women in the United States, and approximately 140,000 women globally succumb to ovarian cancer each year [7]. Research examining the association between ovarian cancer and polycystic ovary syndrome (PCOS) has been conducted, with a notable study by Schildkraut et al. [8] reporting a 2.5-fold risk of ovarian cancer in women with PCOS, which increased to a 10.5-fold risk in those not using oral contraceptives. In contrast, most studies exploring the correlation between PCOS and ovarian cancer have not been able to establish a clear link. An exception is a study by Rossing et al. [9], which investigated connections among infertility, ovarian cancer, and ovulation-inducing drugs. This research suggested a 2.3-fold risk of ovarian cancer associated with clomiphene treatment in these women, a finding supported by subsequent studies [10].

Examining the molecular mechanisms of PCOS that elevate the susceptibility to endometrial and ovarian cancers can provide insights into the pathogenesis of these conditions through bioinformatics data analysis. This study aims to identify the differentially expressed genes (DEGs), pathways, and protein networks shared between Polycystic Ovary Syndrome (PCOS) and endometrial/ovarian cancers. Additionally, it seeks to identify potential targeted therapeutic approaches by assessing common genetic factors identified in PCOS and endometrial/ovarian cancer. The findings may open up new avenues for therapeutic strategies, offering innovative possibilities for treatment.

## MATERIALS AND METHODS

### 1. Data Retrieval

We obtained microarray datasets from the National Center for Biotechnology Information (NCBI) Gene Expression Omnibus to investigate the impact of PCOS and its genetic associations with endometrial and ovarian cancer. The Gene Expression Omnibus (GEO) is a comprehensive and freely accessible database (www.ncbi.nlm.nih.gov/geo) that offers gene expression profiles for various disorders [11].

For our study, we analyzed three distinct microarray datasets from the Gene Expression Omnibus (GEO) database hosted by the NCBI. These datasets corresponded to PCOS, endometrial cancer, and ovarian cancer, identified by the accession numbers GSE199225[12], GSE215413[13], and GSE174670[14], respectively. Our selection criteria included considering only complete samples categorized as either cases or controls. Non-human datasets were excluded, and exclusively human data was utilized.

### 2. Identification of Shared Differentially Expressed Genes (DEGs) in PCOS and Endometrial/Ovarian Cancer

To pinpoint shared DEGs between PCOS and endometrial/ovarian cancer, the online tool GEO2R (www.ncbi.nlm.nih.gov/geo/geo2r/) was employed for group comparison and analysis, offering a user-friendly interface for differential expression analysis [15]. GEO2R facilitates the comparison of gene expression across different experimental conditions within a GEO Series. It integrates DESeq2 for RNA-seq data and GEOquery with limma for microarray data, providing a comprehensive platform for researchers to conduct differential expression analysis and visualize results with high-throughput genomic data.

The datasets GSE199225, GSE215413, and GSE174670 were subjected to GEO2R analysis to identify DEGs, defined as genes with |log2 fold change (log2FC)|>1.0 and an adjusted P-value<0.05. GEO2R not only presents results in a table of genes ordered by P-value but also offers graphic plots for visualizing differentially expressed genes and assessing data set quality. Volcano plots of DEGs from each dataset were obtained through GEO2R analysis, offering a visual representation of statistical significance (-log10 P value) versus the magnitude of change (log2 fold change).

Common DEGs between GSE199225 and GSE215413, as well as GSE199225 and GSE174670 datasets, were identified using GEO2R and visualized using the Venny v2.1 web tool [16]. These common DEGs were considered potential genes associated with the risk of endometrial and ovarian cancer in women with PCOS.

### 3. Analysis of protein-protein interactions (PPI) networks and the identification of hub genes

The evaluation of protein-protein interaction (PPI) networks is fundamental in cellular biology for comprehending protein function and the workings of cellular machinery. The Search Tool for the Retrieval of Interacting Genes (STRING, http://string-db.org) is a database designed for the study of PPI networks, incorporating both physical and functional interactions [17]. In our study, we utilized STRING to construct the PPI network of shared DEGs, focusing on interactions with a score exceeding 0.4. The resulting network was visualized using Cytoscape (Version 3.9.1) [18].

The hub genes of this study were identified using CytoHubba77, a plug-in of Cytoscape.[19]

Hub genes were selected mainly based on their Maximal Clique Centrality (MCC) algorithm, which indicates essentiality of nodes in biological network.

### 4. Functional enrichment analysis

Enrichment analysis of hub genes involved the utilization of the Enrichr web-based tool [20]. This tool was employed for Gene Ontology (GO) and Kyoto Encyclopaedia of Genes and Genomes (KEGG) enrichment analysis, aiming to elucidate the biological functions and signaling pathways associated with the identified hub genes. The analysis encompassed gene ontologies related to biological processes, cellular components, and molecular functions. Additionally, for the exploration of signaling pathways, databases such as Reactome (2022), KEGG pathways (2021), and Wiki Pathways (2023) were investigated.

To establish significance, a statistical threshold criterion of a p-value < 0.05 was applied for the selection of enriched GO terms and pathways.

### 5. Prediction of candidate drugs

The DSigDB database, comprising 19,531 genes and 17,389 compounds, serves as a valuable resource facilitating a direct link between genes and drugs, particularly in the context of drug development studies and translational research [21]. Accessible through the Enrichr webserve (https://amp.pharm.mssm.edu/Enrichr/), the DSigDB database is utilized for analyzing the relationship between drugs and potential targets. In our study, hub genes were submitted to the database to identify potential drug molecules targeting these genes. Subsequently, compounds were ranked based on the adjusted p-value (p < 0.05) and the combined score, calculated using the p-value and z-score by assessing the deviation from the expected rank [22].

## RESULT

### 1. Identification of DEGs and common genes

The GSE199225 data set includes 31 normal skeletal muscle biopsy samples and 30 case skeletal muscle biopsy samples of women affected with PCOS. Supplementary Table 1 and Figure 1A show the results of the differential analysis of the GSE199225 data set, including 2195 DEGs, 2044 down-regulated genes, and 151 up-regulated genes. GSE215413 contains 2 control human endometrium cancer cell line and 9 case LIM1-knocked down (LIM1-KD) human endometrium cancer cell lines. Supplementary Table 2 and Figure 1B show the results of the differential analysis of the GSE215413 data set, including 3136 DEGs, 2795 down-regulated genes, and 341 up-regulated genes.

**FIGURE 1.**
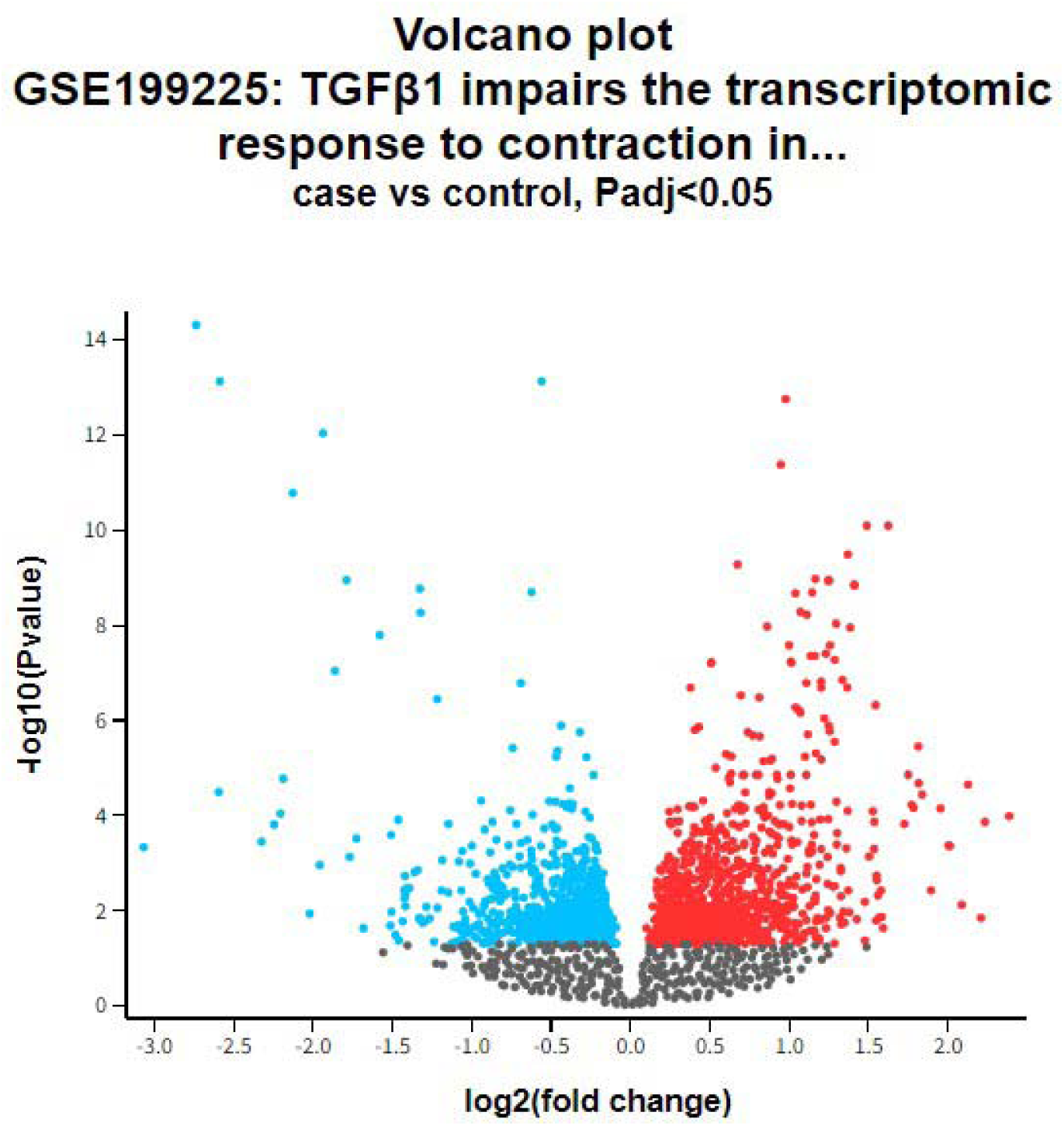
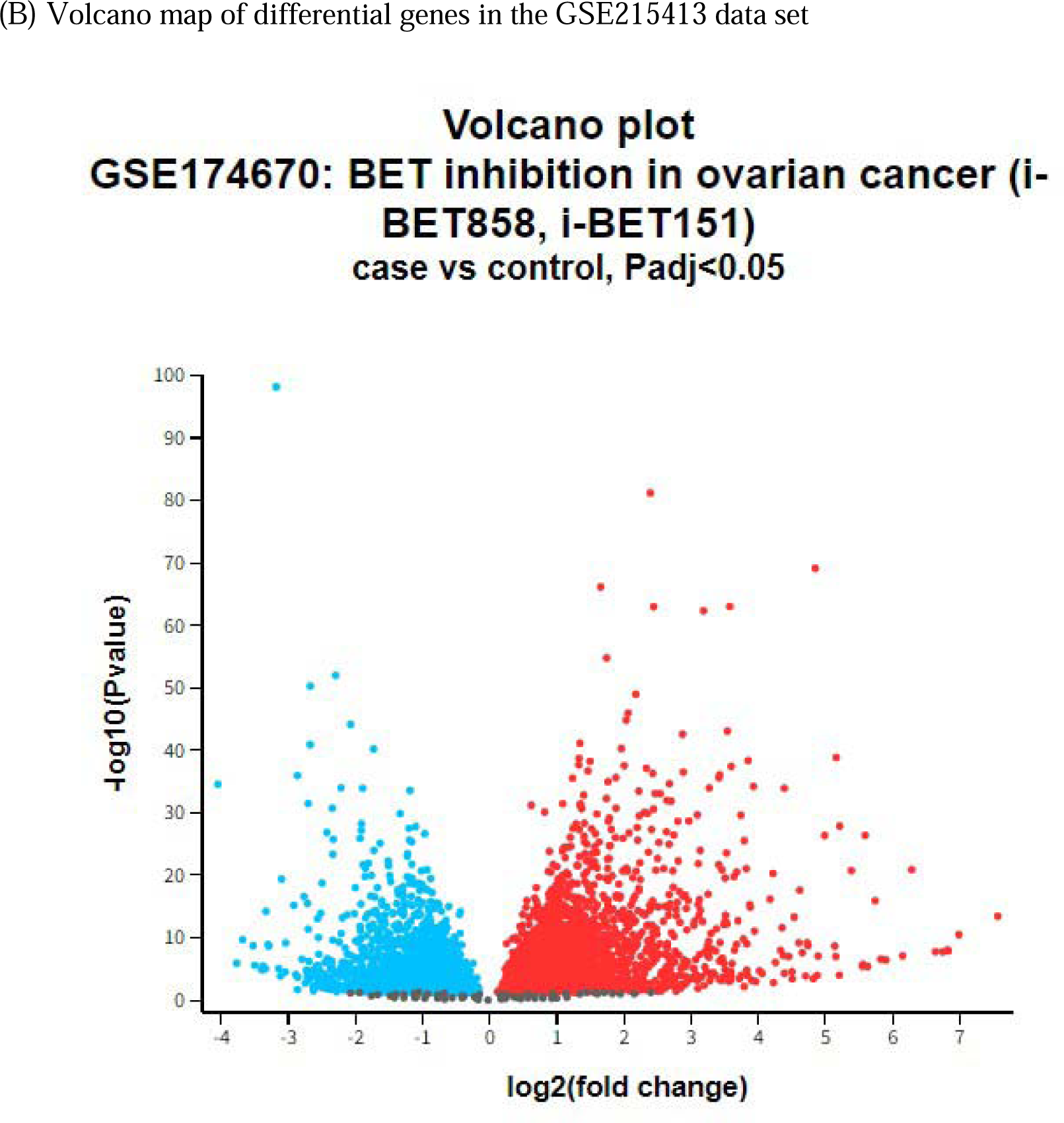
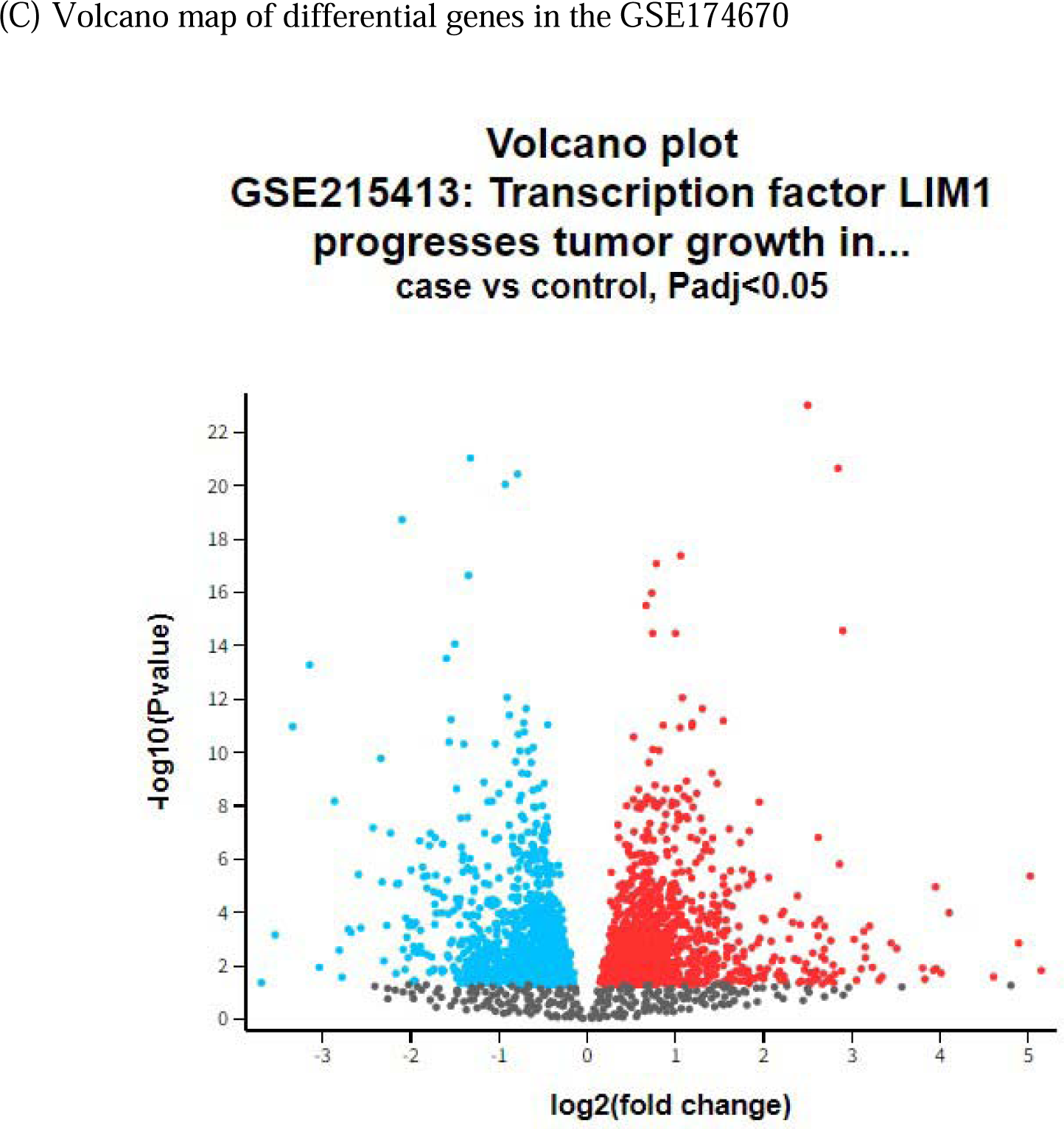
| (A) Volcano map of differential genes in the GSE199225 data set

GSE174670 datasets contains 6 control samples and 12 case samples for the study. Supplementary table 3 figure 1C shows the results of the differential analysis of the GSE174670 data set, including 6724 DEGs, 5415 down-regulated genes, and 1309 up-regulated genes. The detail of the included datasets is showed in table.1

**Table 1:**
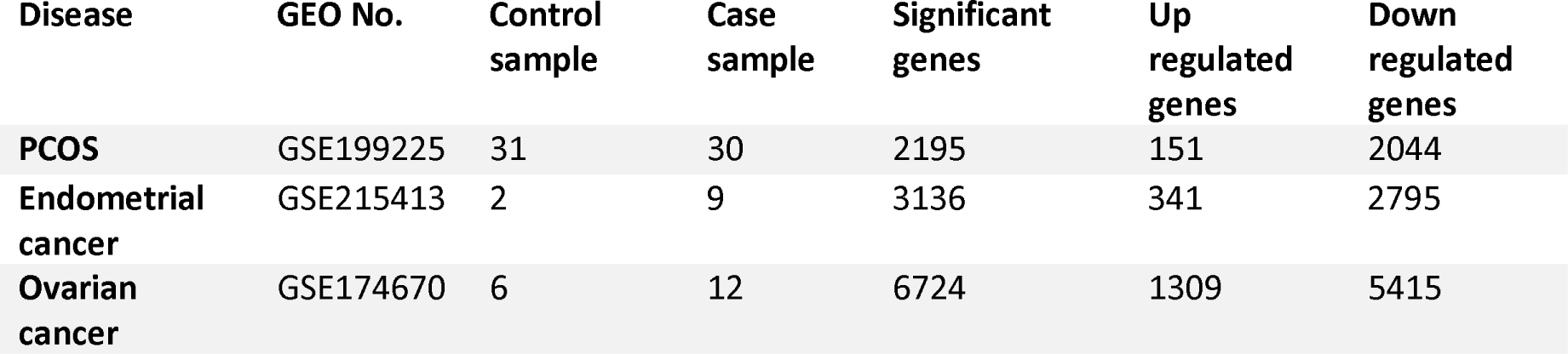
Details information of the selected microarray data

| Disease | GEO No. | Control sample | Case sample | Significant genes | Up regulated genes | Down regulated genes |
| --- | --- | --- | --- | --- | --- | --- |
| **PCOS** | GSE199225 | 31 | 30 | 2195 | 151 | 2044 |
| **Endometrial cancer** | GSE215413 | 2 | 9 | 3136 | 341 | 2795 |
| **Ovarian cancer** | GSE174670 | 6 | 12 | 6724 | 1309 | 5415 |

The Venn diagram visualization revealed 344 genes shared between PCOS and endometrial cancer (Figure 2A), and 716 genes common to both PCOS and ovarian cancer. (Figure 2B)

**Figure 2.**
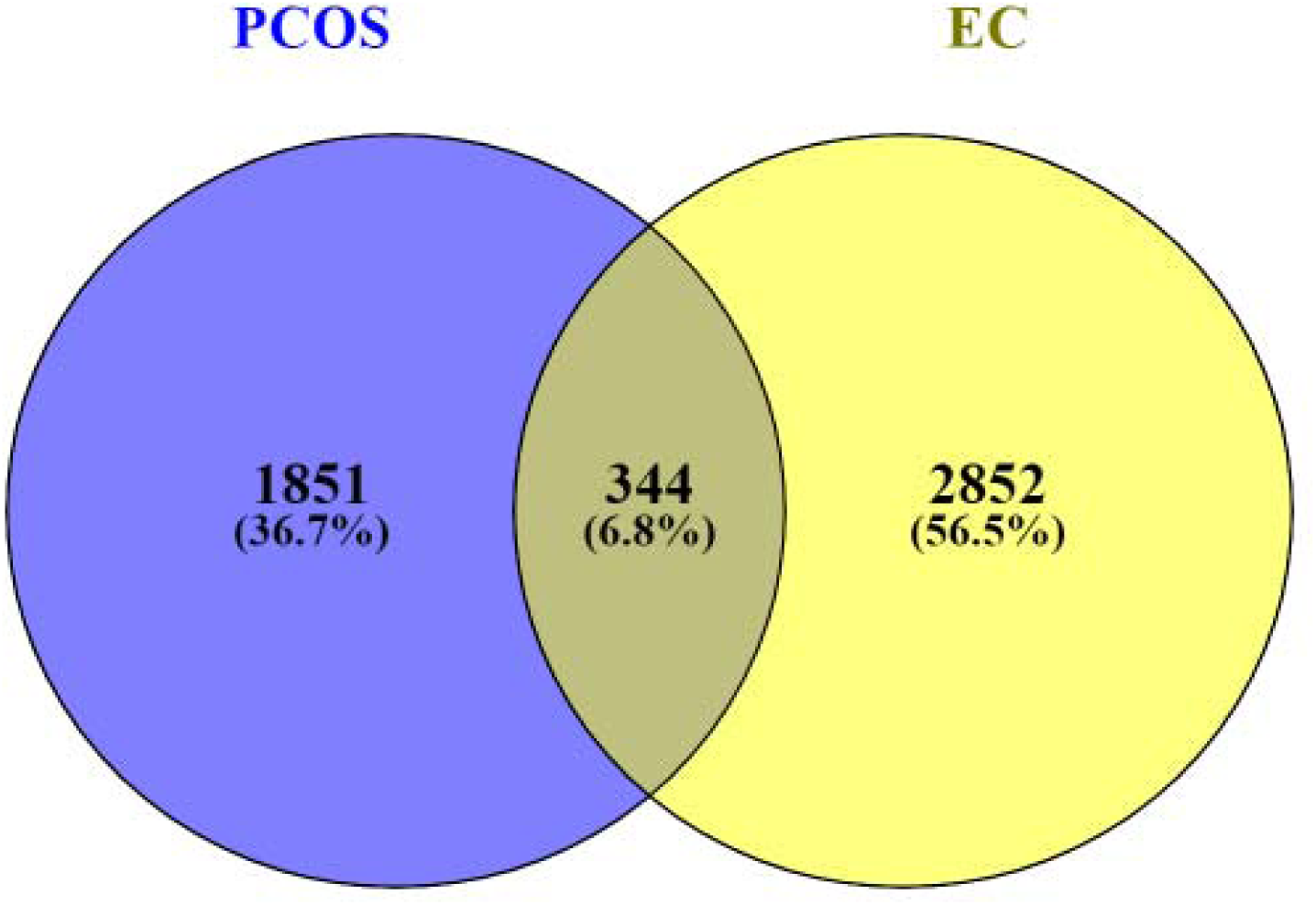
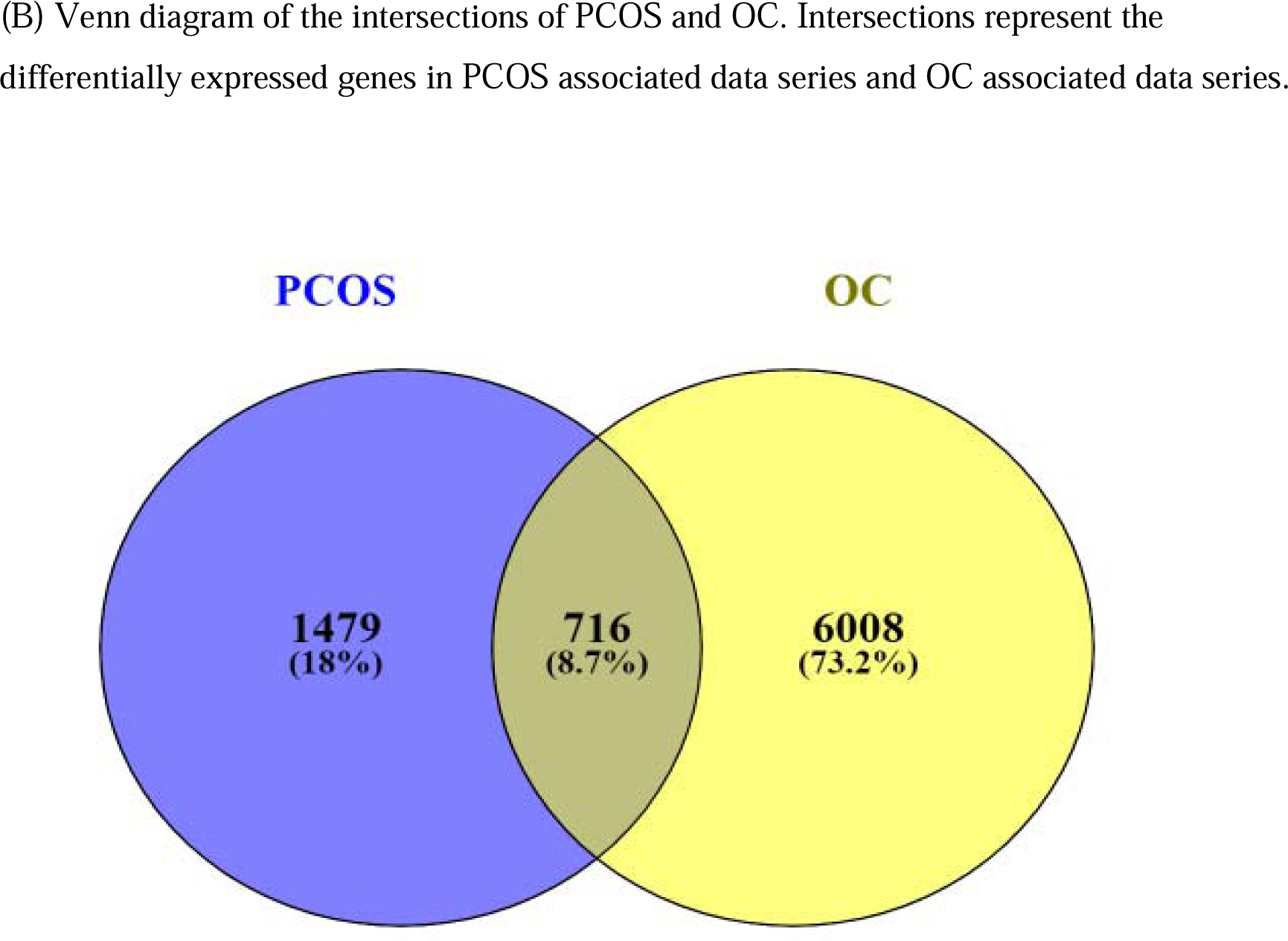
(a)Venn diagram of the intersections of PCOS and EC. Intersections represent the differentially expressed genes in PCOS associated data series and EC associated data series

### 2. Analysis of the Protein-Protein Interaction (PPI) network and the identification of Hub genes

The PPI network analysis of 344 common DEGs between PCOS and endometrial cancer (EC) and 716 common DEGs between PCOS and ovarian cancer (OC) was conducted using the STRING platform, with a medium confidence threshold (score > 0.4) for constructing the PPI network. The results were visualized using Cytoscape software. Subsequently, the top 10 hub genes were identified through the implementation of 11 algorithms from the cytoHubba plug-in in Cytoscape. The maximum centrality clique (MCC) algorithm was specifically employed to identify the top 10 hub nodes, with the ranking visualized using a red-to-yellow colour gradient.

The top 10 hub genes associated with PCOS and endometrial cancer include RECQL4, RAD54L, ATR, CHTF18, WDHD1, CDT1, PLK1, PKMYT1, RAD18, and RPL3 (Table 2.A) (supplementary table 4). The network comprised 10 nodes and 35 edges. (Figure 3.A) Similarly, the top 10 hub genes associated with PCOS and ovarian cancer include HMOX1, TXNRD1, NQO1, GCLC, GSTP1, PRDX1, SOD1, GPX3, BOP1, and BYSL (Table 2.B) (supplementary table 5). The network for this association consisted of 10 nodes and 45 edges. (Figure 3.B)

**Figure 3.**
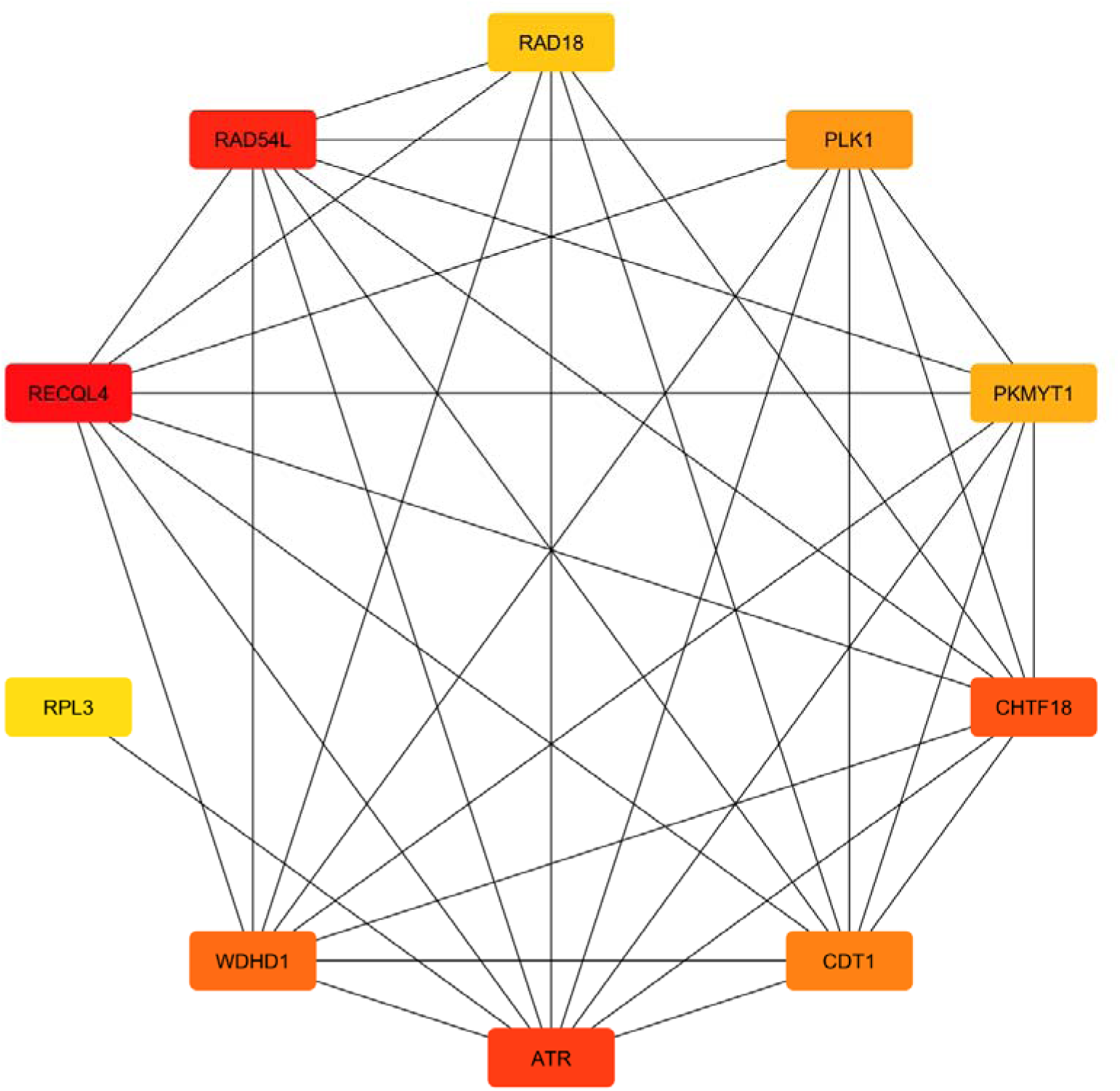
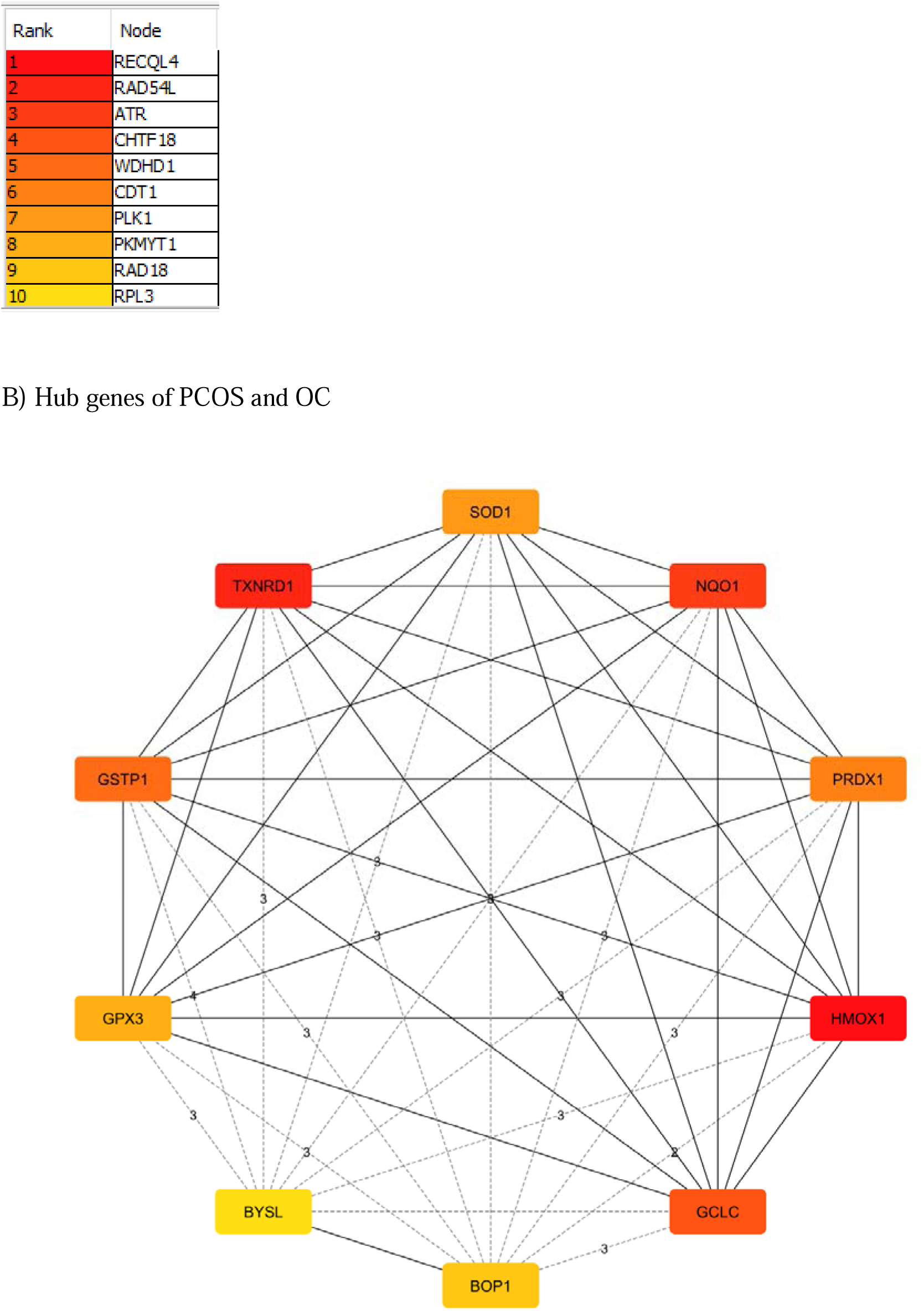
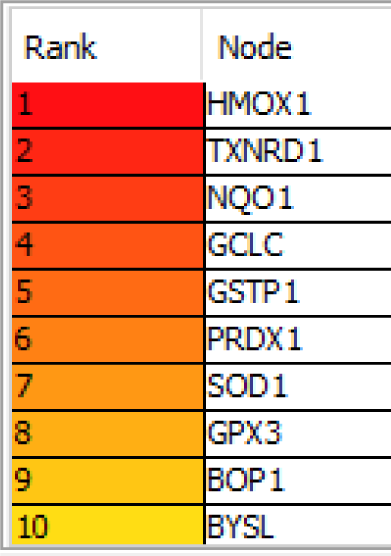
Hub genes screening. The colour of the nodes is illustrated from red to yellow in descending order of MCC score. Gray lines highlight the interactions; line thickness refers to the interaction score provided by STRING. The redder the colour of the gene in the network, the higher the connectivity of the gene with other genes.

**Table 2.** Hub genes identified through PPI analysis. A) PCOS and EC B) PCOS and OC Maximal Clique Centrality (MCC) scores indicated essentiality of the gene in biological network. the greater the value, the more important the gene

| Hub genes | Mcc Score | Description |
| --- | --- | --- |
| RECQL4 | 6840 | RecQ like helicase 4 |
| RAD54L | 6828 | RAD54 like |
| ATR | 6671 | ATR serine/threonine kinase |
| CHTF18 | 6606 | chromosome transmission fidelity factor 18 |
| WDHD1 | 6600 | WD repeat and HMG-box DNA binding protein 1 |
| CDT1 | 6062 | chromatin licensing and DNA replication factor 1 |
| PLK1 | 5262 | polo like kinase 1 |
| PKMYT1 | 5162 | protein kinase, membrane associated tyrosine/threonine 1 |
| RAD18 | 1764 | RAD18 E3 ubiquitin protein ligase |
| RPL3 | 1024 | ribosomal protein L3 |

B)
| Hub gene | MCC Score | Description |
| --- | --- | --- |
| HMOX1 | 5362 | heme oxygenase 1 |
| TXNRD1 | 5347 | thioredoxin reductase 1 |
| NQO1 | 5311 | NAD(P)H quinone dehydrogenase 1 |
| GCLC | 5305 | glutamate-cysteine ligase catalytic subunit |
| GSTP1 | 5194 | glutathione S-transferase pi 1 |
| PRDX1 | 5177 | peroxiredoxin 1 |
| SOD1 | 5113 | superoxide dismutase 1 |
| GPX3 | 5089 | glutathione peroxidase 3 |
| BOP1 | 1725 | BOP1 ribosomal biogenesis factor |
| BYSL | 1720 | bystin like |

### 3. Pathway and functional enrichment analysis

A diverse array of signaling pathways and Gene Ontology (GO) terms play a crucial role in the orchestration and development of diseases, especially in complex disorders. To analyze the GO and pathway enrichment of the Hub genes, we utilized the Enrichr online tool. The significance of terms was determined based on the P-value. The GO analysis comprised three categories: biological process (2023), cellular component (2023), and molecular function (2023). The top 10 significant terms from each category were summarized in Table (3), (4) and presented as bar graphs in Figure (4)(5).

**Table 3.** Ontological analysis of Hub genes between PCOS and EC

| Category | GO ID | Term | P- values | Genes |
| --- | --- | --- | --- | --- |
| GO<br>Biological process | GO:0006259 | DNA Metabolic Process | 0.000008259 | RECQL4, WDHD1, RAD18, ATR |
|  | GO:0006281 | DNA Repair | 0.000008604 | RECQL4, WDHD1, RAD18, ATR |
|  | GO:0006974 | DNA Damage Response | 0.00002564 | RECQL4, WDHD1, RAD18, ATR |
|  | GO:0000086 | G2/M Transition of Mitotic Cell Cycle | 0.0001567 | PLK1, PKMYT1 |
|  | GO:0044839 | Cell Cycle G2/M Phase Transition | 0.0001737 | PLK1, PKMYT1 |
|  | GO:0034502 | Protein Localization to Chromosome | 0.0001737 | PLK1, ATR |
|  | GO:0031570 | DNA Integrity Checkpoint Signaling | 0.0001917 | CDT1, ATR |
|  | GO:0032200 | Telomere Organization | 0.0004323 | RECQL4, ATR |
| GO<br>Cellular component | GO:0043232 | Intracellular Non-Membrane-Bounded Organelle | 0.0001477 | RECQL4, RPL3, PLK1, PKMYT1, ATR |
|  | GO:0005634 | Nucleus | 0.001782 | RECQL4, CDT1, RPL3, PLK1, CHTF18, RAD18, ATR |
|  | GO:0005694 | Chromosome | 0.002880 | RECQL4, ATR |
|  | GO:0043231 | Intracellular Membrane-Bounded Organelle | 0.004322 | RECQL4, CDT1, RPL3, PLK1, CHTF18, RAD18, ATR |
|  | GO:0042405 | Nuclear Inclusion Body | 0.004990 | RAD18 |
|  | GO:0022625 | Cytosolic Large Ribosomal Subunit | 0.02570 | RPL3 |
|  | GO:0015934 | Large Ribosomal Subunit | 0.02570 | RPL3 |
|  | GO:0005876 | Spindle Microtubule | 0.03349 | PLK1 |
| GO<br>Molecular function | GO:0000217 | DNA Secondary Structure Binding | 0.0001178 | RECQL4, RAD18 |
|  | GO:0004674 | Protein Serine/Threonine Kinase Activity | 0.0005440 | PLK1, PKMYT1, ATR |
|  | GO:0009378 | Four-Way Junction Helicase Activity | 0.002498 | RECQL4 |
|  | GO:0140666 | Annealing Activity | 0.002498 | RECQL4 |
|  | GO:0032356 | Oxidized DNA Binding | 0.003495 | RECQL4 |
|  | GO:0000405 | Bubble DNA Binding | 0.003495 | RECQL4 |
|  | GO:0032357 | Oxidized Purine DNA Binding | 0.004492 | RECQL4 |
|  | GO:0043138 | 3'-5' DNA Helicase Activity | 0.006979 | RECQL4 |

**Table 4.**
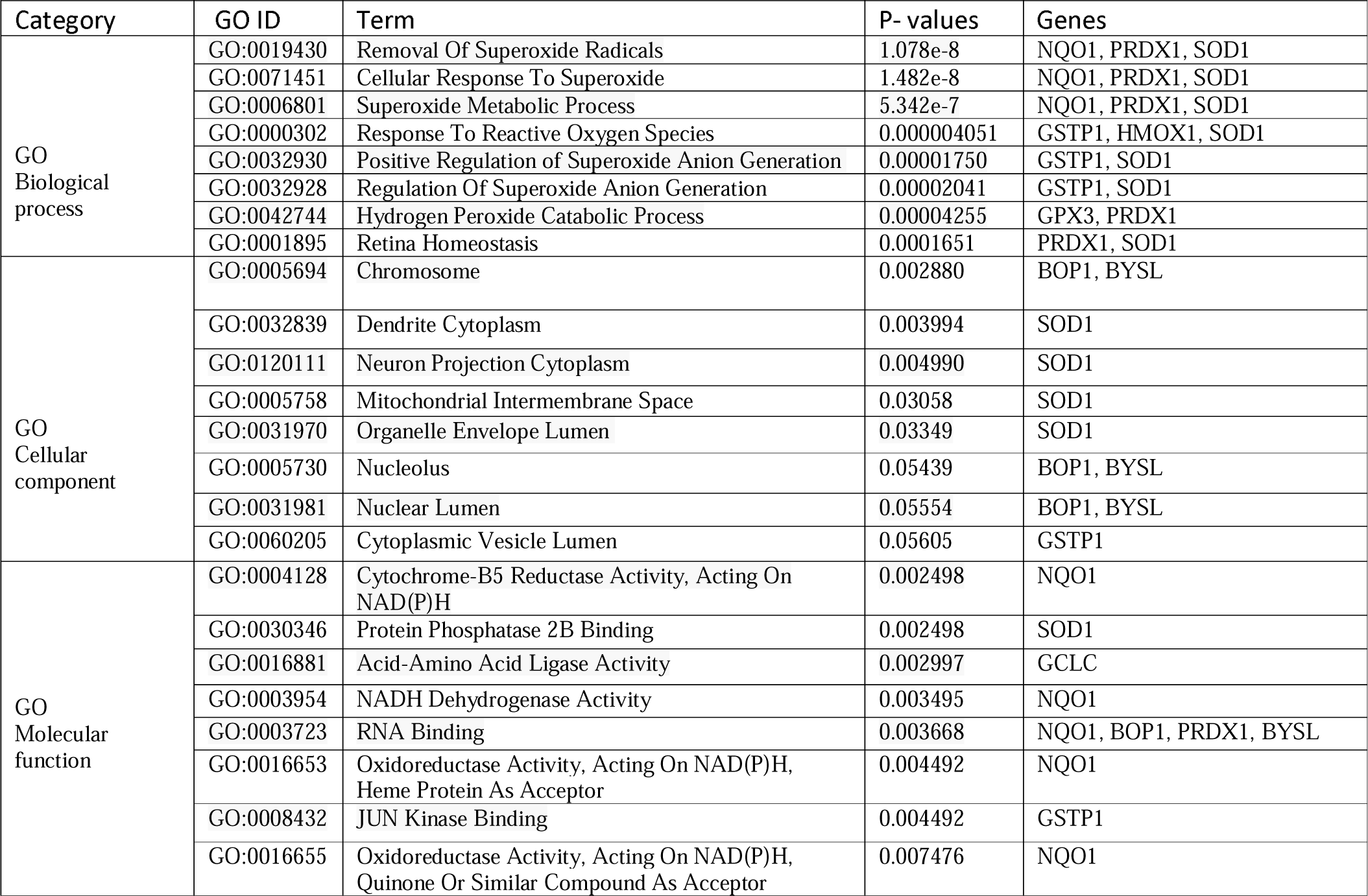
Ontological analysis of Hub genes between PCOS and OC

| Category | GO ID | Term | P- values | Genes |
| --- | --- | --- | --- | --- |
| GO Biological process | GO:0019430 | Removal Of Superoxide Radicals | 1.078e-8 | NQO1, PRDX1, SOD1 |
|  | GO:0071451 | Cellular Response To Superoxide | 1.482e-8 | NQO1, PRDX1, SOD1 |
|  | GO:0006801 | Superoxide Metabolic Process | 5.342e-7 | NQO1, PRDX1, SOD1 |
|  | GO:0000302 | Response To Reactive Oxygen Species | 0.000004051 | GSTP1, HMOX1, SOD1 |
|  | GO:0032930 | Positive Regulation Of Superoxide Anion Generation | 0.00001750 | GSTP1, SOD1 |
|  | GO:0032928 | Regulation Of Superoxide Anion Generation | 0.00002041 | GSTP1, SOD1 |
|  | GO:0042744 | Hydrogen Peroxide Catabolic Process | 0.00004255 | GPX3, PRDX1 |
|  | GO:0001895 | Retina Homeostasis | 0.0001651 | PRDX1, SOD1 |
| GO Cellular component | GO:0005694 | Chromosome | 0.002880 | BOP1, BYSL |
|  | GO:0032839 | Dendrite Cytoplasm | 0.003994 | SOD1 |
|  | GO:0120111 | Neuron Projection Cytoplasm | 0.004990 | SOD1 |
|  | GO:0005758 | Mitochondrial Intermembrane Space | 0.03058 | SOD1 |
|  | GO:0031970 | Organelle Envelope Lumen | 0.03349 | SOD1 |
|  | GO:0005730 | Nucleolus | 0.05439 | BOP1, BYSL |
|  | GO:0031981 | Nuclear Lumen | 0.05554 | BOP1, BYSL |
|  | GO:0060205 | Cytoplasmic Vesicle Lumen | 0.05605 | GSTP1 |
| GO Molecular function | GO:0004128 | Cytochrome-B5 Reductase Activity, Acting On NAD(P)H | 0.002498 | NQO1 |
|  | GO:0030346 | Protein Phosphatase 2B Binding | 0.002498 | SOD1 |
|  | GO:0016881 | Acid-Amino Acid Ligase Activity | 0.002997 | GCLC |
|  | GO:0003954 | NADH Dehydrogenase Activity | 0.003495 | NQO1 |
|  | GO:0003723 | RNA Binding | 0.003668 | NQO1, BOP1, PRDX1, BYSL |
|  | GO:0016653 | Oxidoreductase Activity, Acting On NAD(P)H, Heme Protein As Acceptor | 0.004492 | NQO1 |
|  | GO:0008432 | JUN Kinase Binding | 0.004492 | GSTP1 |
|  | GO:0016655 | Oxidoreductase Activity, Acting On NAD(P)H, Quinone Or Similar Compound As Acceptor | 0.007476 | NQO1 |

**Figure 4.**
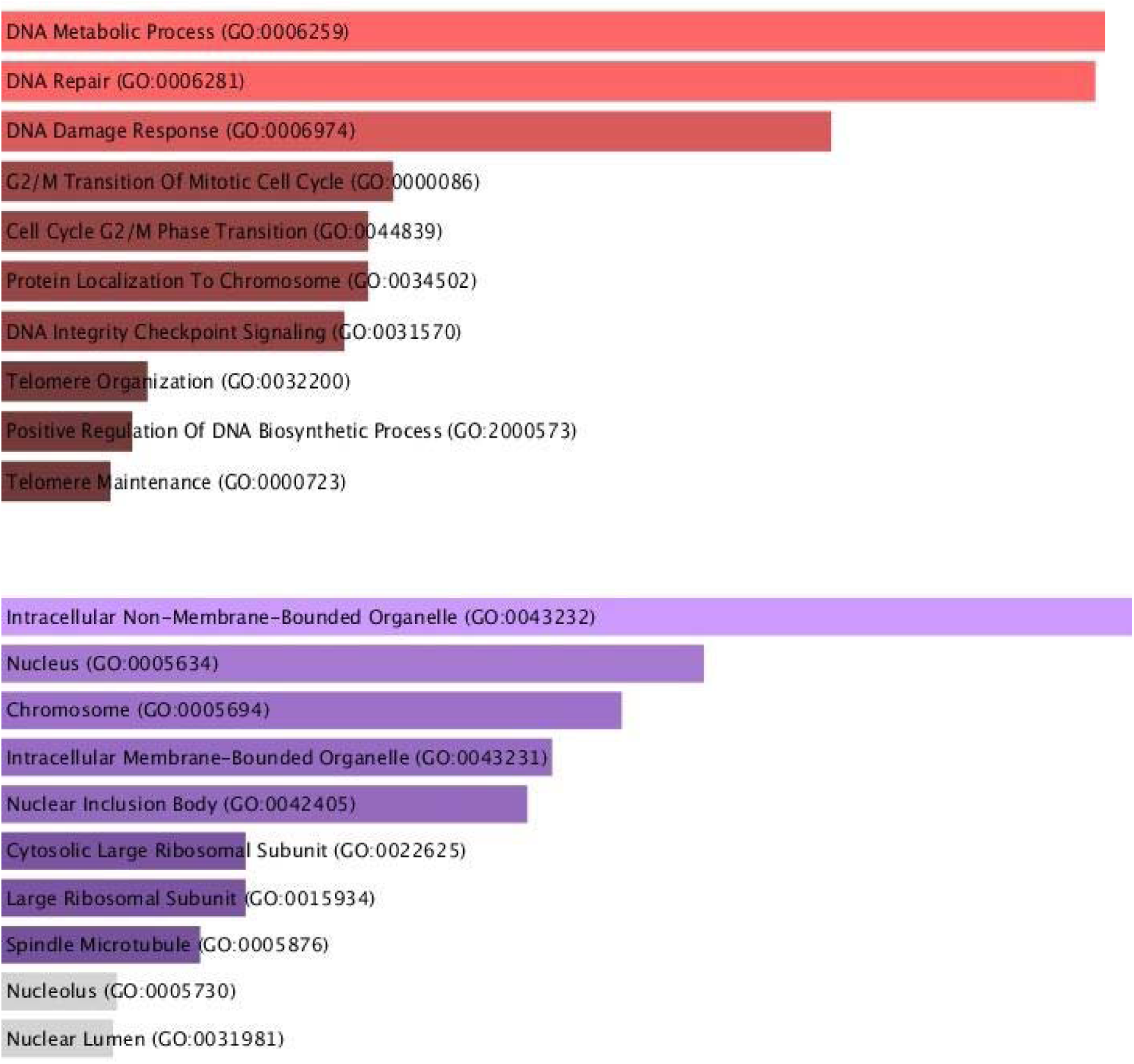
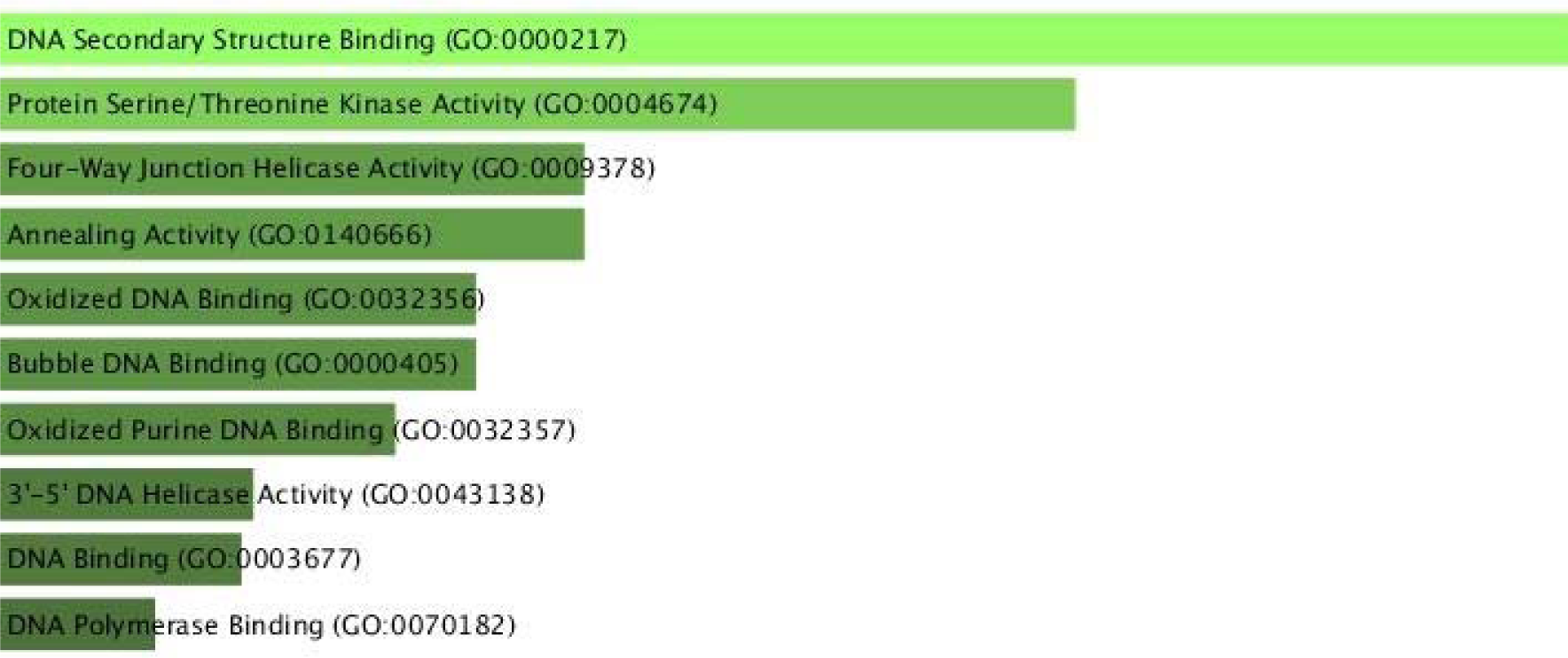
GO terms of Hub genes between PCOS and EC(A) Biological Processes, (B) cellular component, (C) molecular function

**Figure 5.**
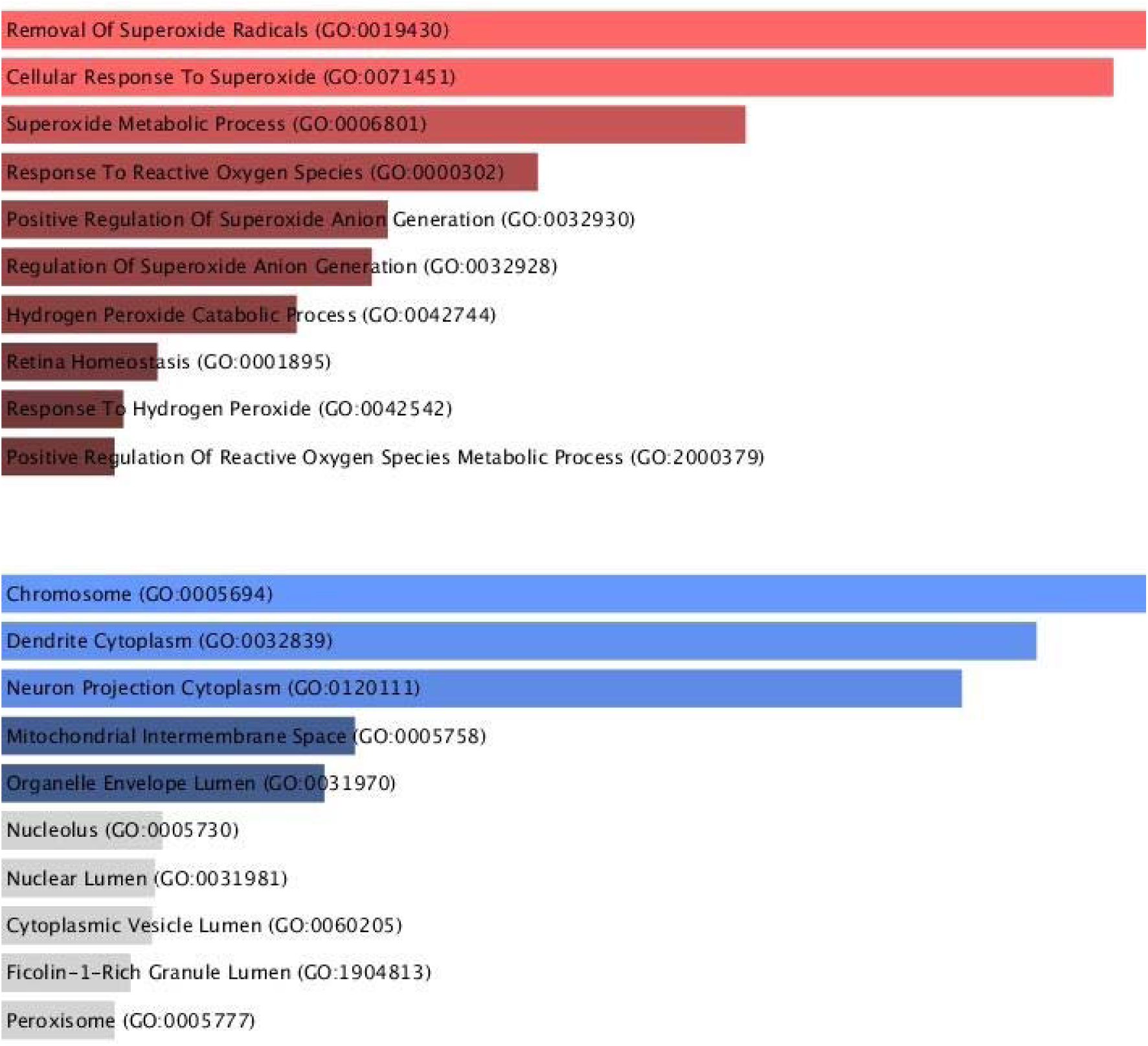
GO terms of Hub genes between PCOS and OC(A) Biological Processes, (B) cellular component, (C) molecular function

**Figure 6.**
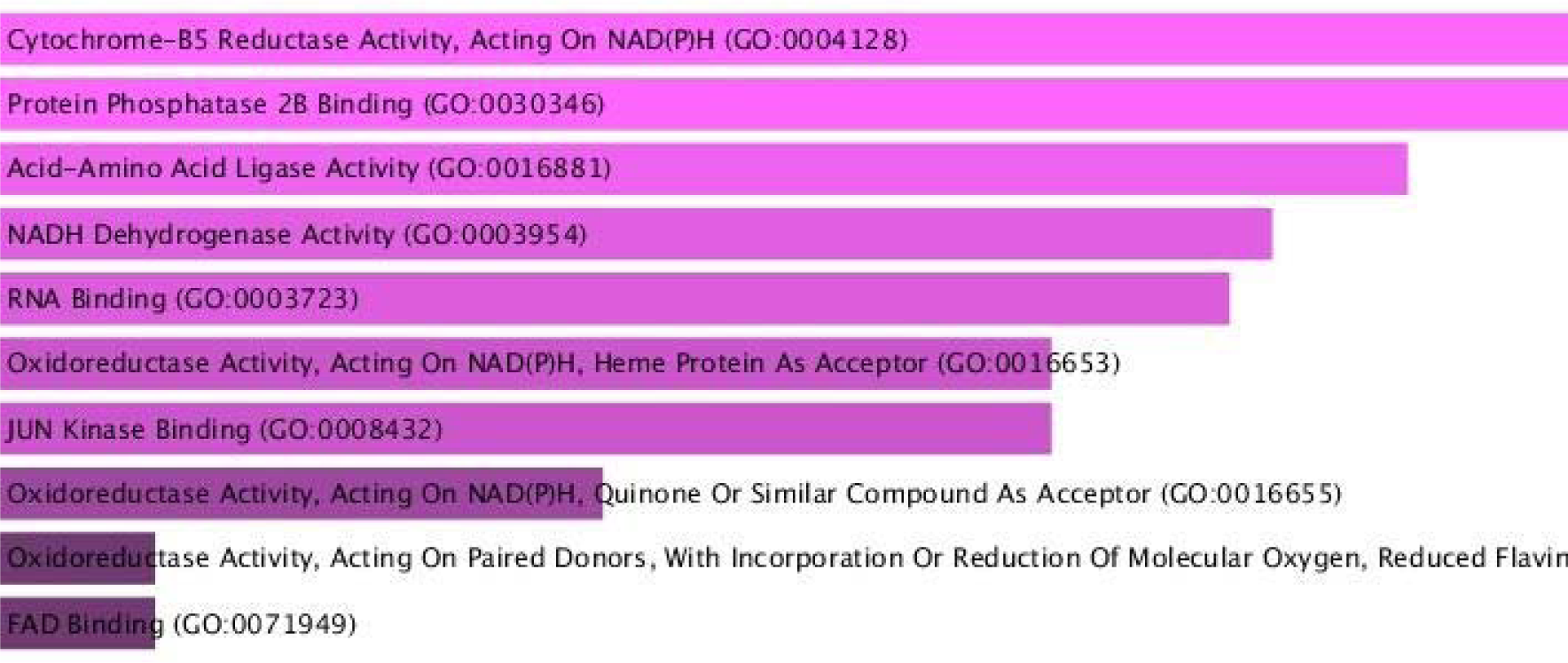
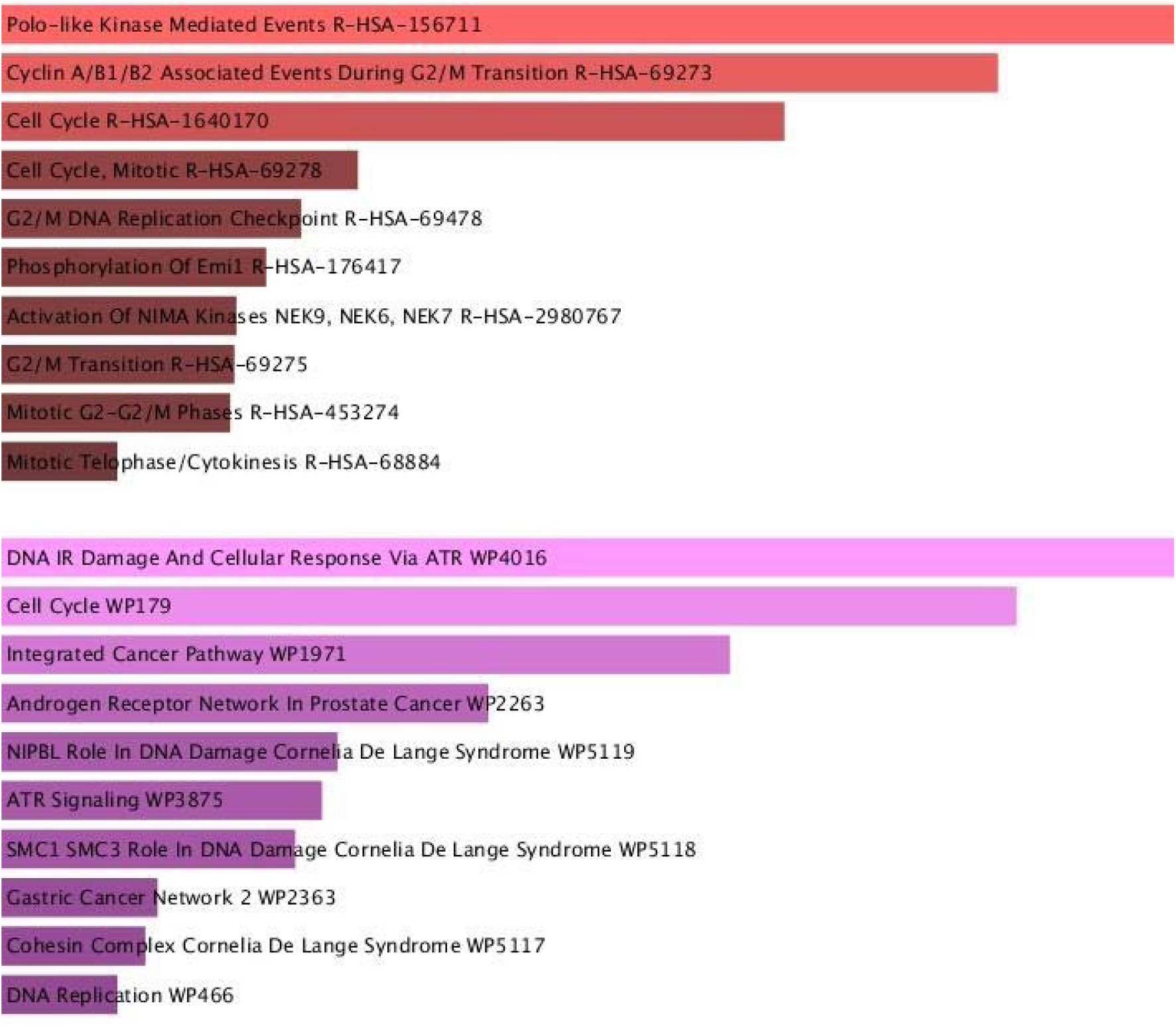
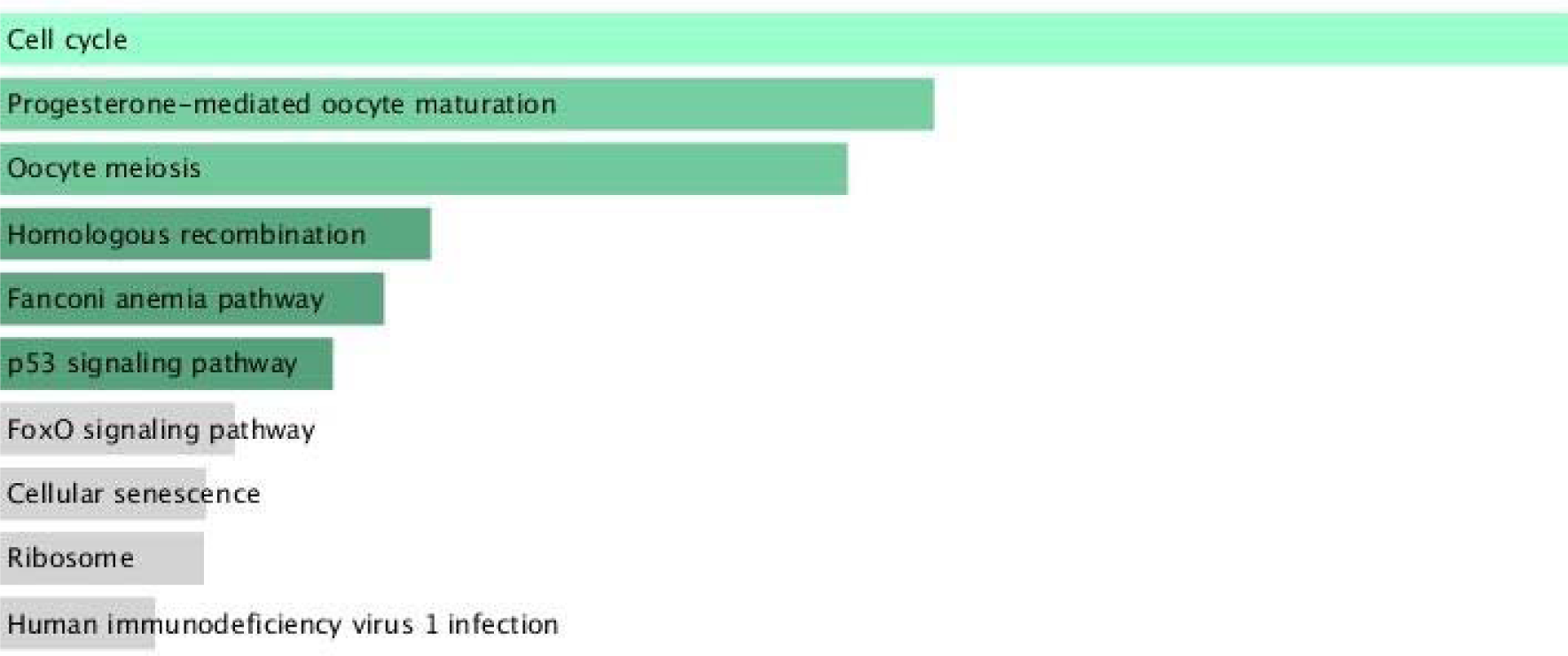
Pathway enrichment analysis of Hub genes between PCOS and EC. (A) Reactome Pathway, (B) Wikipathway, (C) KEGG Human Pathway

**Figure 7.**
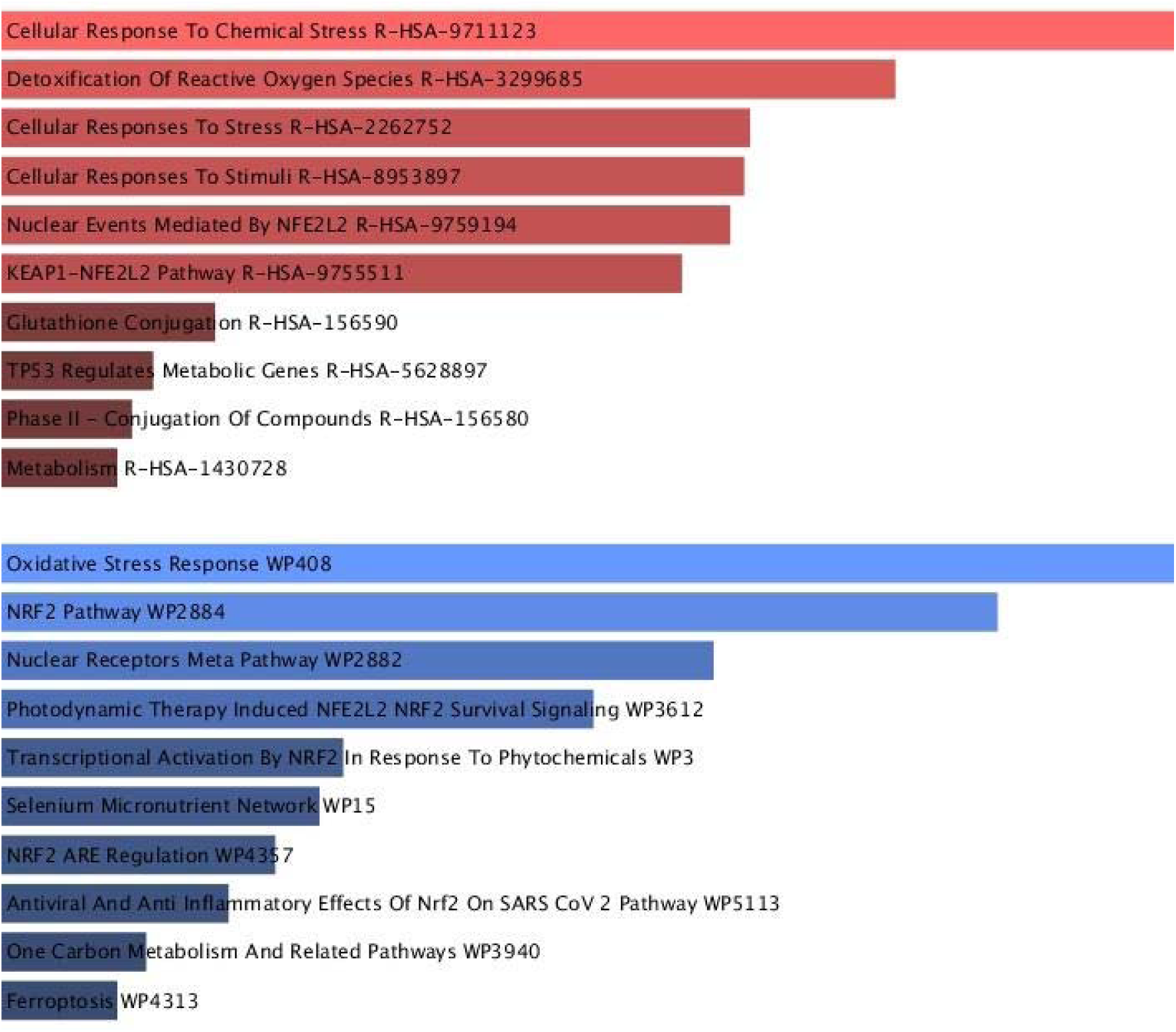
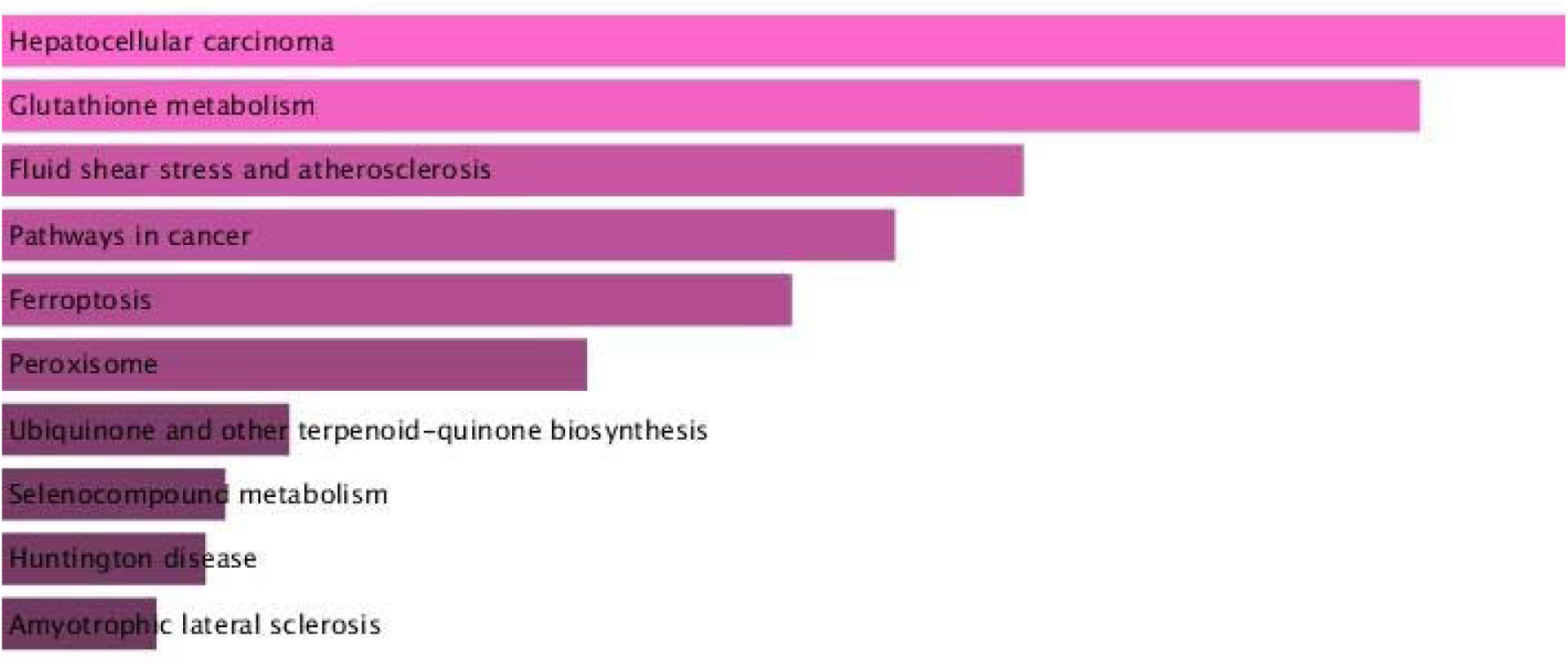
Pathway enrichment analysis of Hub genes between PCOS and EC. (A) Reactome Pathway, (B) Wikipathway, (C) KEGG Human Pathway

This analysis helped identify key pathways and gene ontologies that could establish connections between PCOS and endometrial/ovarian cancer using hub genes.

### 4. Prediction of drug candidate

Utilizing the Enrichr platform, which is based on the DSigDB database, we identified and ranked the top 10 potential therapeutic compounds based on their P-values in relation to hub genes. These compounds were considered as promising pharmacological targets for PCOS and endometrial cancer (EC) (Table 7) and PCOS and ovarian cancer (OC) (Table 8).

**Table 5.**
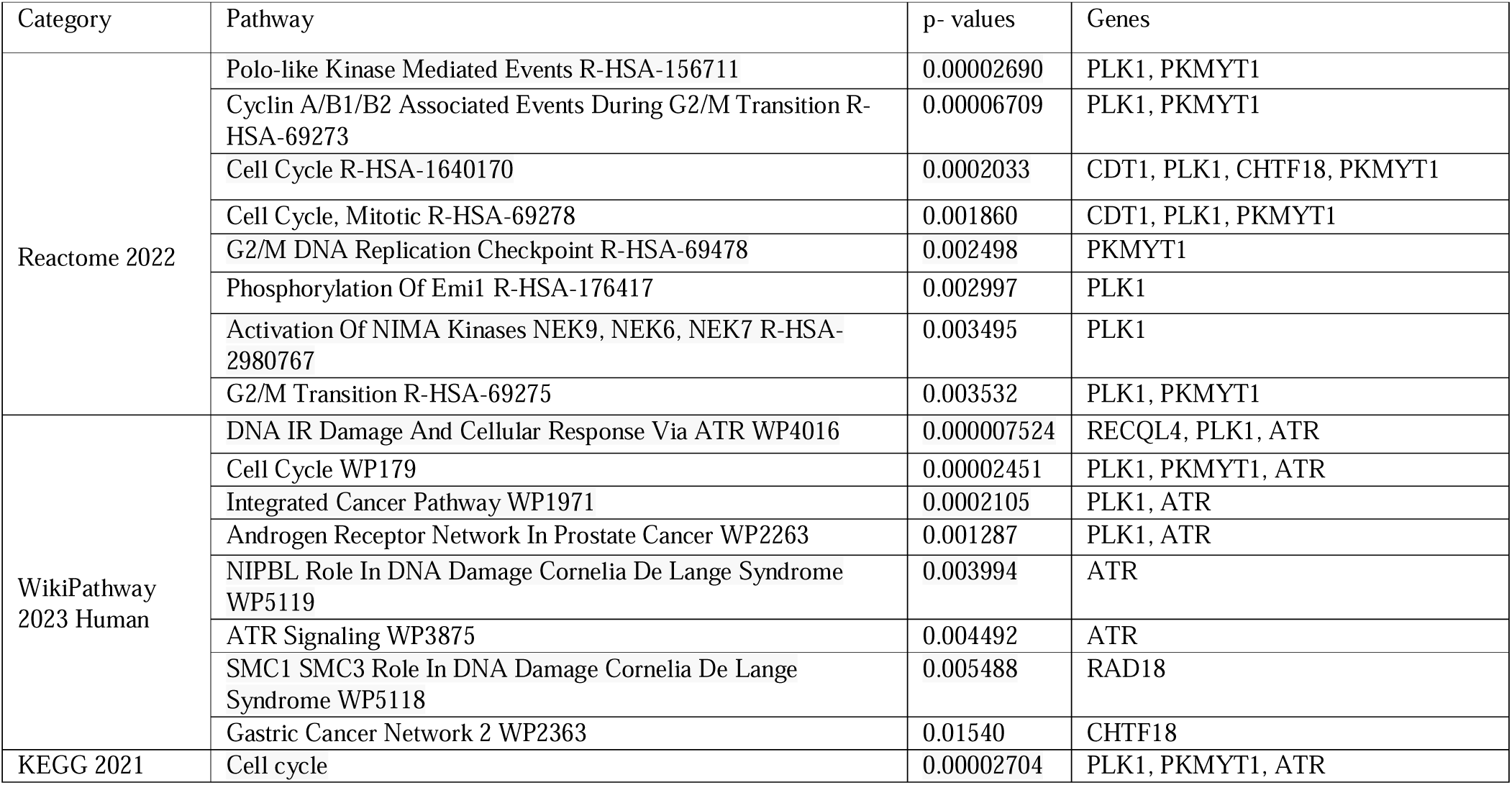
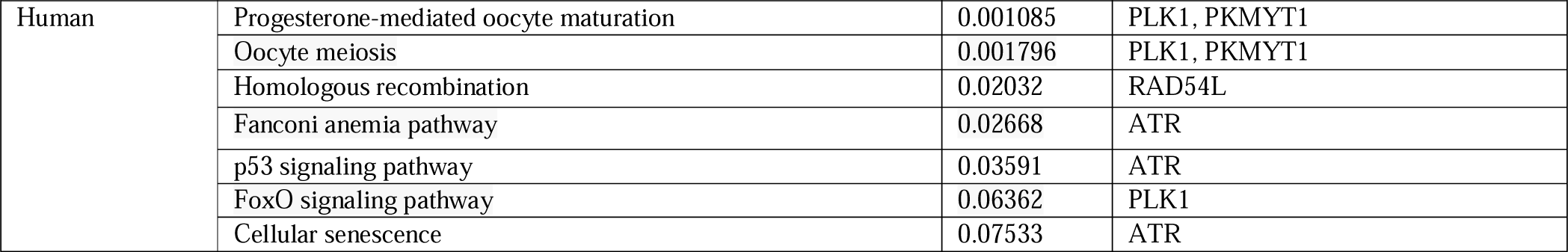
Pathway enrichment analysis of Hub genes between PCOS and EC

**Table 6.**
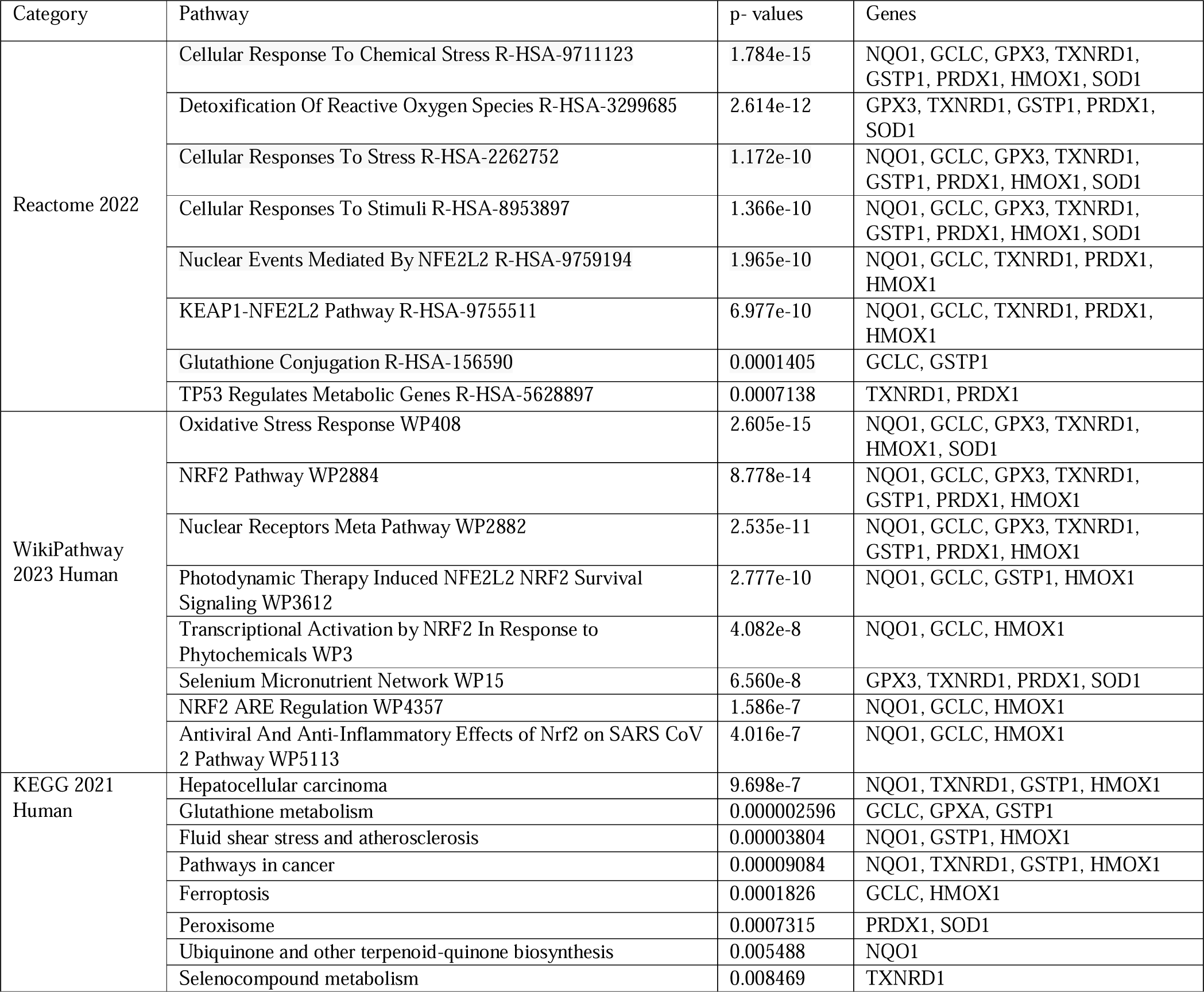
Pathway enrichment analysis of Hub genes between PCOS and OC

**Table 7.** Prediction of top 10 candidate drugs for PCOS and EC

| Drug name | P-value | Genes |
| --- | --- | --- |
| testosterone CTD 00006844 | 8.984e-9 | RECQL4, WDHD1, CDT1, PLK1, RAD54L, CHTF18, PKMYT1, RAD18 |
| troglitazone CTD 00002415 | 2.183e-7 | RECQL4, CDT1, PLK1, RAD54L, CHTF18, PKMYT1, |
| calcitriol CTD 00005558 | 3.127e-7 | RECQL4, WDHD1, CDT1, PLK1, RAD54L, CHTF18, PKMYT1, RAD18 |
| Enterolactone CTD 00001393 | 0.000002289 | RECQL4, WDHD1, CDT1, PLK1, RAD54L, PKMYT1 |
| COUMESTROL CTD 00005717 | 0.000004649 | RECQL4, WDHD1, CDT1, PLK1, RAD54L, PKMYT1, RAD18 |
| 0297417-0002B PC3 DOWN | 0.000005272 | WDHD1, CDT1, PKMYT1 |
| quercetin CTD 00006679 | 0.00001277 | RECQL4, WDHD1, CDT1, PLK1, RAD54L, CHTF18, PKMYT1, ATR |
| resveratrol CTD 00002483 | 0.00004129 | RECQL4, WDHD1, CDT1, PLK1, RAD54L, ATR |
| LITHOCHOLIC ACID BOSS | 0.00007266 | CDT1, PLK1 |
| Prazosin hydrochloride BOSS | 0.0001327 | CDT1, RAD54L |

**Table 8.** Prediction of top 10 candidate drugs for PCOS and OC

| Drug name | P-value | Genes |
| --- | --- | --- |
| alpha-Hexylcinnamaldehyde CTD 00002937 | 2.605e-15 | NQO1, GCLC, GPX3, TXNRD1, HMOX1, SOD1 |
| cinnamyl alcohol CTD 00001006 | 4.579e-15 | NQO1, GCLC, GPX3, TXNRD1, HMOX1, SOD1 |
| Bandrowski's base CTD 00002216 | 9.018e-15 | NQO1, GCLC, GPX3, TXNRD1, HMOX1, SOD1 |
| 1-chloro-2,4-dinitrobenzene CTD 00005848 | 1.803e-14 | NQO1, GCLC, GPX3, TXNRD1, GSTP1, PRDX1, HMOX1, SOD1 |
| CHLOROBENZENE CTD 00001495 | 4.118e-14 | GPX3, TXNRD1, GSTP1, HMOX1, SOD1 |
| CHEBI:18224 CTD 00001728 | 6.053e-14 | NQO1, GCLC, GPX3, TXNRD1, HMOX1, SOD1 |
| eugenol CTD 00005949 | 2.258e-13 | NQO1, GCLC, GPX3, TXNRD1, GSTP1, HMOX1, SOD1 |
| MANEB CTD 00006239 | 9.244e-13 | NQO1, GCLC, TXNRD1, PRDX1, SOD1 |
| disulfiram CTD 00005860 | 6.882e-12 | NQO1, GCLC, GPX3, TXNRD1, HMOX1, SOD1 |
| tert-Butylhydroquinone CTD 00000961 | 7.979e-12 | NQO1, GCLC, TXNRD1, GSTP1, HMOX1 |

Notably, three pharmacological molecules—testosterone (CTD 00006844), calcitriol (CTD 00005558), and quercetin (CTD 00006679)—were found to interact with the majority of genes in the PCOS and EC context. In the case of PCOS and OC, the two drugs that interacted with most of the genes were 1-chloro-2,4-dinitrobenzene (CTD 00005848) and eugenol (CTD 00005949).

## DISCUSSION

Several studies have suggested an elevated risk of endometrial and ovarian cancer in women with polycystic ovary syndrome (PCOS), indicating a potential link between PCOS, endometrial cancer (EC), and ovarian cancer (OC). However, the underlying mechanisms remain unclear.

In this study, a series of bioinformatics analyses were conducted to identify hub genes associated with PCOS and EC/OC. A total of 10 hub genes were identified from the differentially expressed genes (DEGs) in PCOS and EC, including RECQL4, RAD54L, ATR, CHTF18, WDHD1, CDT1, PLK1, PKMYT1, RAD18, and RPL3. Similarly, another set of 10 hub genes emerged from the DEGs in PCOS and OC, including HMOX1, TXNRD1, NQO1, GCLC, GSTP1, PRDX1, SOD1, GPX3, BOP1, and BYSL.

Gene Ontology (GO) analysis was performed using the 20 hub genes identified across the three datasets of PCOS, EC, and OC. Noteworthy biological processes in the GO analysis of PCOS and EC included DNA Metabolic Process, DNA Repair, and DNA Damage Response. Conversely, in the GO analysis of PCOS and OC, significant processes included Removal of Superoxide Radicals, Cellular Response to Superoxide, and Superoxide Metabolic Process.

DNA damage response and repair pathways are implicated in endometrial cancer development [23]. Increased expression of obesity-associated genes in women with PCOS may contribute to endometrial cancer progression [24]. PCOS, marked by endocrine and metabolic disruptions, often accompanies oxidative stress. Addressing internal oxidative stress could be a potential therapeutic strategy [25]. Reactive oxygen species (ROS) play a key role in signaling, but excessive or prolonged ROS production is linked to cancer initiation and progression [26]. Oxidative stress is implicated in the onset of ovarian cancer [27]. Superoxide dismutases (SODs), enzymes responsible for neutralizing superoxide free radicals, play vital roles in cancer cell development, growth, and survival [28].

KEGG pathway enrichment analysis unveiled that the most significant pathways in the context of PCOS and endometrial cancer include Cell cycle, Progesterone-mediated oocyte maturation, and Oocyte meiosis. Dysregulation of key cell cycle regulators, like cyclin B1, cyclin D1, cyclin E, p16, p21, p27, p53, and cdk2, is crucial in endometrial cancer progression [29]. Progesterone, vital for oocyte maturation and embryo development, has significant effects in vitro and in vivo, with conflicting results [30]. Acting through PR-A and PR-B, progesterone plays a crucial role in uterine physiology, regulating development, implantation, and acting as a tumor suppressor in endometrial cancer cells [31]. Elevated mRNA instability index (mRNAsi) in uterine corpus endometrial carcinoma correlates with reduced overall survival, possibly involving oocyte meiosis and cell cycle pathways [32].

In the case of PCOS and ovarian cancer, the top three significant pathways identified through KEGG pathway enrichment analysis are Hepatocellular carcinoma, Glutathione metabolism, and Fluid shear stress and atherosclerosis. PCOS, marked by chronic anovulation and hyperandrogenism, is closely linked to obesity and insulin resistance (IR). These are key factors in non-alcoholic steatohepatitis (NASH), which can lead to cirrhosis with potential complications like portal hypertension, liver failure, and hepatocellular carcinoma [33]. Glutathione metabolism plays a dual role in cancer, offering both protection and promoting pathology. Elevated glutathione levels may shield tumor cells, conferring resistance to chemotherapy [34]. Women with PCOS display increased serum levels of homocysteine, glutathione, lipid peroxidation, and superoxide, potentially affecting reproductive health and raising cancer risk [35]. Fluid shear stress activates the Ras-MEKK-JNK pathway, impacting endothelial gene expression in atherosclerosis-related diseases [36]. Disruptions in this pathway could potentially lead to cancer.

The hub genes selected from the protein-protein interaction (PPI) network of PCOS and endometrial cancer include RECQL4, RAD54L, ATR, CHTF18, WDHD1, CDT1, PLK1, PKMYT1, RAD18, and RPL3. Ribosomal protein S3 (RPS3) negatively modulates RECQL4’s unwinding activity, influencing active DNA repair during cellular stress. RECQL4, crucial for preventing tumorigenesis and maintaining genome integrity, is upregulated in cancer [37]. RAD54L, involved in homologous recombination and DNA repair, is markedly upregulated, correlating with a worse prognosis in multiple cancers [38]. ATR, monitoring DNA structure, phosphorylates proteins in DNA damage response pathways. Truncating mutations in ATR are identified in endometrioid endometrial cancer, producing a shortened or incomplete ATR protein [39].

Based on the protein-protein interaction (PPI) network analysis of PCOS and ovarian cancer, hub genes were identified, including HMOX1, TXNRD1, NQO1, GCLC, GSTP1, PRDX1, SOD1, GPX3, BOP1, and BYSL.Heme oxygenase-1 (HMOX1) is a vital enzyme in heme breakdown. In endometriotic lesions, where endometrial tissue grows outside the uterus, macrophages infiltrate and play a crucial role in processing heme released during menstruation. HMOX1, also called HO-1, processes heme within macrophages and is elevated in Endometriosis-associated ovarian cancer [41]. Thioredoxin reductase 1 (TXNRD1) is a key element of the thioredoxin system, a significant cellular antioxidant defense. It aids in reducing thioredoxin, essential for neutralizing reactive oxygen species (ROS) and sustaining proteins in a functional state. Elevated metabolic activity in cancer cells results in increased oxidative stress, leading to the upregulation of TXNRD1 in various cancer types [42].

The identified hub genes from the analysis were used to predict potential drugs using the DSigDB database. The top three chemical molecules for the association between polycystic ovary syndrome (PCOS) and endometrial cancer (EC) were testosterone (CTD 00006844), calcitriol (CTD 00005558), and quercetin (CTD 00006679). For the association between PCOS and ovarian cancer (OC), the top drugs were 1-chloro-2,4-dinitrobenzene (CTD 00005848) and eugenol (CTD 00005949). Testosterone may contribute to endometrial cancer via membrane-initiated signaling pathways. Steroid inhibitors could reduce hormone-dependent endometrial cancer [43]. Calcitriol, the active vitamin D form, may have anti-tumor effects through the VDR gene mechanism [44]. Quercetin inhibits endometrial fibrosis, offering potential in treatments and prevention [45].

For the PCOS and OC association, 1-chloro-2,4-dinitrobenzene (DNCB) is linked to assessing biological risk in primary lung cancer [46]. Eugenol inhibits breast precancerous lesion progression via the HER2/PI3K-AKT pathway, showing promise for breast cancer treatment [47]. Cisplatin combined with eugenol effectively restrains ovarian cancer growth, reducing side population cells and suppressing cancer stem cell resistance through the Notch-Hes1 pathway [48].

Since our findings were obtained by pure bioinformatics analysis, so the function of hub genes and prospective medicines is required to be further confirmed by scientific investigation in vitro and in vivo.

## CONCLUSION

In Conclusion, this study presents compelling evidence supporting the link between PCOS and Endometrial/Ovarian cancer, with common DEGs identified through the screening of genome expression datasets. Bioinformatics analysis revealed gene signatures and regulatory patterns. Additionally, we elucidated various molecular and gene ontology pathways, offering a clear perspective on the genetic connection between EC/OC and PCOS. The gene expression study highlighted ten hub genes, most of which significantly contribute to both PCOS and EC/OCr. Furthermore, we delved into the molecular mechanisms underlying these signatures and identified potential small drug molecules associated with the hub genes. However, given the data-driven nature of our research, it is crucial to conduct further experimental and clinical studies to validate the identified molecular signatures and potential drugs.

## Notes

### Competing Interest Statement

The authors have declared no competing interest.

